# Season-specific responses of freshwater ciliate communities to top-down and bottom-up experimental manipulations

**DOI:** 10.1101/2025.03.06.641887

**Authors:** Usman Asghar, Indranil Mukherjee, Bettina Sonntag, Caio César Pires de Paula, Vojtěch Kasalický, Paul-Adrian Bulzu, Anusha Priya Singh, Tanja Shabarova, Kasia Piwosz, Karel Šimek

**Affiliations:** Biology Centre of the Czech Academy of Sciences, Institute of Hydrobiology, Na Sádkách 7, 37005, České Budějovice, Czech Republic; Faculty of Science, University of South Bohemia, 37005, České Budějovice, Czech Republic; Research Department for Limnology, Mondsee, Universität Innsbruck, A-5310 Mondsee, Austria; National Marine Fisheries Research Institute, ul. Kołłątaja 1, 81-332 Gdynia, Poland

**Keywords:** Ciliates, aquatic food web, microbial loop, experimental manipulations, freshwater reservoir, quantitative protargol staining, long-read amplicon sequencing

## Abstract

In aquatic microbial food webs, ciliates represent an important trophic link in the energy transfer from prokaryotes, algae, and heterotrophic nanoflagellates (HNF) to higher trophic levels. However, the trophic role of abundant small ciliates (< 20 µm) is not clearly understood. To unveil their trophic linkages, we conducted two experiments manipulating both top-down and bottom-up controlling factors, thus modulating the trophic cascading and bacterial prey availability for protists during contrasting spring and summer seasons with samples collected from a freshwater meso-eutrophic reservoir. Water samples were size fractionated, to modify food web complexity, i.e. 10-µm, 20-µm and unfiltered control and amended with bacterial prey additions. The samples were analyzed by morphological and sequencing techniques. The bacterial amendments triggered strong ciliate growth following the peaks of HNF in the 10-µm and 20-µm treatments, reflecting a trophic cascading from HNF to raptorial prostome ciliates (*Balanion planctonicum* and *Urotricha* spp.) in spring. In summer, HNF and ciliates peaked simultaneously, suggesting the important trophic cascade from bacteria to bacterivorous scuticociliates (*Cyclidium glaucoma* and *Cinetochilum margaritaceum*) and HNF. In spring, unfiltered treatments showed stronger ciliate top-down control by zooplankton than in summer. The sequence analysis revealed season-specific manipulation-induced shifts in ciliate communities and their large cryptic diversity. However, morphological and molecular analyses also revealed considerable discrepancies in the abundance of major ciliate taxa. The ciliate communities responded to our experimental manipulations in season-specific fashion, thereby highlighting the different roles of ciliates as an intermediate trophic link between prokaryotes and higher trophic levels.

**IMPORTANCE:** Ciliates are an important trophic link in aquatic microbial food webs. In this study, we used the food web manipulation techniques to reveal their complex trophic interactions during seasonally different plankton scenarios occurring in spring and summer. Manipulating top-down controlling factors (grazing pressure of micro- and metazooplankton grazers) and bottom-up factors (an availability of bacterial prey) shaped distinctly the complexity and dynamics of natural plankton communities and thus yielded significant changes in ciliate community dynamics. The experimentally simplified plankton and ciliate communities responded to our manipulations in season-specific fashions, reflected in different roles of ciliates as an intermediate trophic link between prokaryotes and higher trophic levels. This study also demonstrates that the combination of morphological and molecular analyses is essential for providing robust and ecologically meaningful results due to the reliability in quantifying the major ciliate taxa and their trophic role.

## INTRODUCTION

Ciliates are an extremely diverse and ubiquitous group of unicellular eukaryotes (1, 2). The concept of microbial loop (3), attributed to the important roles of ciliates in the aquatic microbial food webs, as they contribute significantly to the trophic complexity by linking different trophic level of the food webs (4–7). For instance, ciliates possess various feeding modes allowing them to graze on detritus, prokaryotes, unicellular eukaryotes, other ciliates and diatoms and are preyed upon by meta-zooplankton, fish larvae and planktivorous fish (8, 9).

Considering temporal microbial community dynamics, changes in availability of food resources, community structure of zooplankton and controlling physical factors (e.g., solar irradiation, temperature, etc.), the recurrent seasonal patterns in temperate freshwater ecosystems were set to the following four different phases: spring phytoplankton bloom, clearwater phase, summer/autumn and winter (10, 11). This plankton seasonality is also reflected (5, 12–15) in recurring successional patterns in both community composition and abundance of planktonic ciliates in temperate lakes. According to the description of the revised Plankton Ecology Group (PEG) model (11), and of other recent studies (16, 17), ciliated protists are the first and the most voracious grazers of phytoplankton during spring bloom events. The ciliate community during spring is dominated by medium-sized (<30 µm) algivorous oligotrichs and small (>20 µm) prostomatids (*Balanion planctonicum* and *Urotricha* spp.), which are usually replaced first by omnivorous/mixotrophic ciliates and then by small bacterivorous scuticociliates (18).

Additionally, the ciliate community dynamics and abundance are also influenced by the trophic status of respective lakes. In eutrophic lakes, the peak in the abundance of ciliates generally occurs during the spring phytoplankton bloom and again in late summer (10, 14, 18–21). In mesotrophic and meso-eutrophic lakes, the abundance of ciliates reaches its annual maximum in spring (10, 18). In contrast, in eutrophic lakes the seasonal maxima are recorded in late summer since in spring ciliates experience a very strong top-down control by zooplankton predators (5).

Generally, the ciliate assemblage in spring consists mainly of larger herbivorous ciliate species of the class Spirotrichea (*Pelagostrombidium* spp., *Tintinnidium fluviatile* and *Codonella cratera*) (20), whose dominance contributes significantly to the total ciliate biomass. Especially in mesotrophic and meso-eutrophic systems, small algivorous Prostomatea (e.g., *B. planctonicum*) show a steep increase in abundance after a bloom of flagellated algae and contribute about 60-80% to the total ciliate count during their spring maximum (16, 22, 23). The spring maximum of algivorous ciliates is usually followed by the succession of several mixotrophic ciliates (*Pelagohalteria viridis*, *Histiobalantium bodamicum*, *Limnostrombidium viride, Askenasia* spp. and *Mesodinium* spp.) and heterotrophic algivorous *Urotricha* spp. (10, 18), while omnivorous and bacterivorous ciliate species (*Halteria grandinella* and *Rimostrombidium* spp., *Cyclidium* spp., *Uronema* spp.) prevail during a summer/autumn period (20, 24).

Both food availability (bottom-up) and predation by larger zooplankton (top-down) have been suggested as the main factors controlling the abundance and community composition of ciliates in freshwater lakes (7, 13, 25). However, most of the previous studies have analyzed the effects of top-down and bottom-up processes in highly complex natural food webs, where numerous other confounding factors such as solar irradiation, temperature, pH, oxygen, predation, food quality, etc. influences the structuring of natural plankton communities (11). This considerably limits our ability to distinguish the true causal relationships and to refine our knowledge on the trophic role of the key members of planktonic ciliates.

In this study, we examined the responses of natural ciliate communities, originating from the contrasting spring and summer plankton phases in Římov reservoir, Czech Republic to experimental manipulations substantially modulating both top-down and bottom-up controlling factors. The plankton samples from the seasonally distinct phases were subjected to the same experimental manipulations, in which we significantly increased bacterial prey availability for protistan communities and, in parallel, we simplified a food web structure and trophic interactions by size fractionation (10-µm, 20-µm and unfiltered) of the plankton samples (Figure 1). This resulted in different levels of food web complexity brought about by simplifying the complex natural zooplankton communities as top predators in the unfiltered fraction in contrast to considerably simplified interactions of largely protist-dominated communities in the 20-µm and 10-µm fractions.

**Figure 1.**
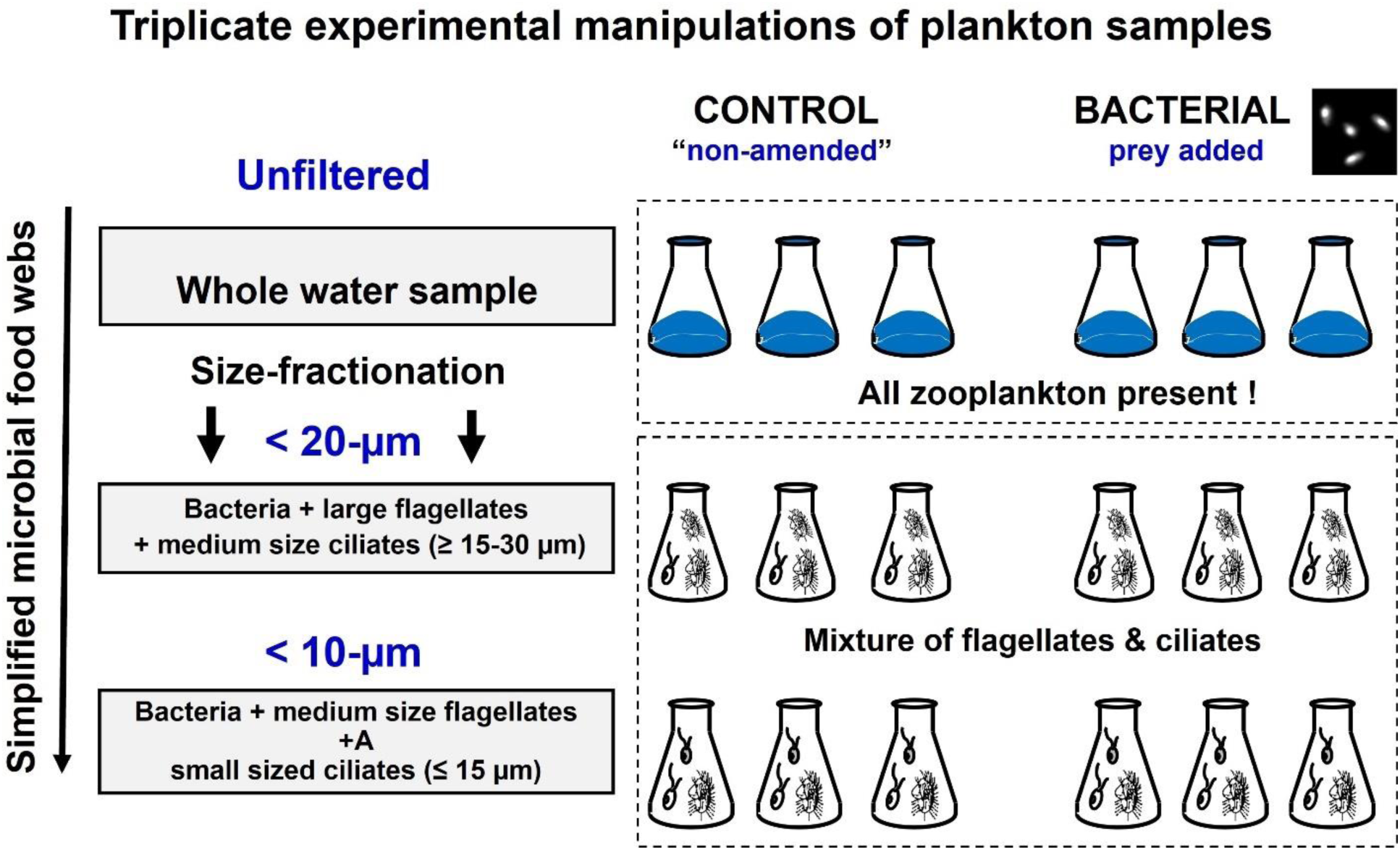
The experimental design showing the treatments: unfiltered - natural microbial and zooplankton communities, 20-μm filtered - supporting growth of micro-sized ciliates and large flagellates and the 10-μm filtered - supporting growth of nano-sized ciliates and small flagellates.

## RESULTS

The environmental parameters from the samples taken from the Římov reservoir in spring and summer are shown in Table 1. In general, based on both quantitative protargol staining method (QPS) and sequencing data, the comparison of ciliate communities showed that the ciliate assemblage was more diverse in summer than in spring. Shannon diversity index also showed significantly higher diversity in the summer (3.36 ± 0.62) as compared to spring samples (2.88 ± 0.54) (Figure S1). We were able to detect 18 species of ciliates in spring and 20 in summer using QPS (Table 2). Taxonomically, we found a greater diversity in the class Spirotrichea in spring (44 % of the total assemblage), while in summer the Oligohymenophorea were more diverse (30 %), followed by Spirotrichea (25 %).

**Table 1.**
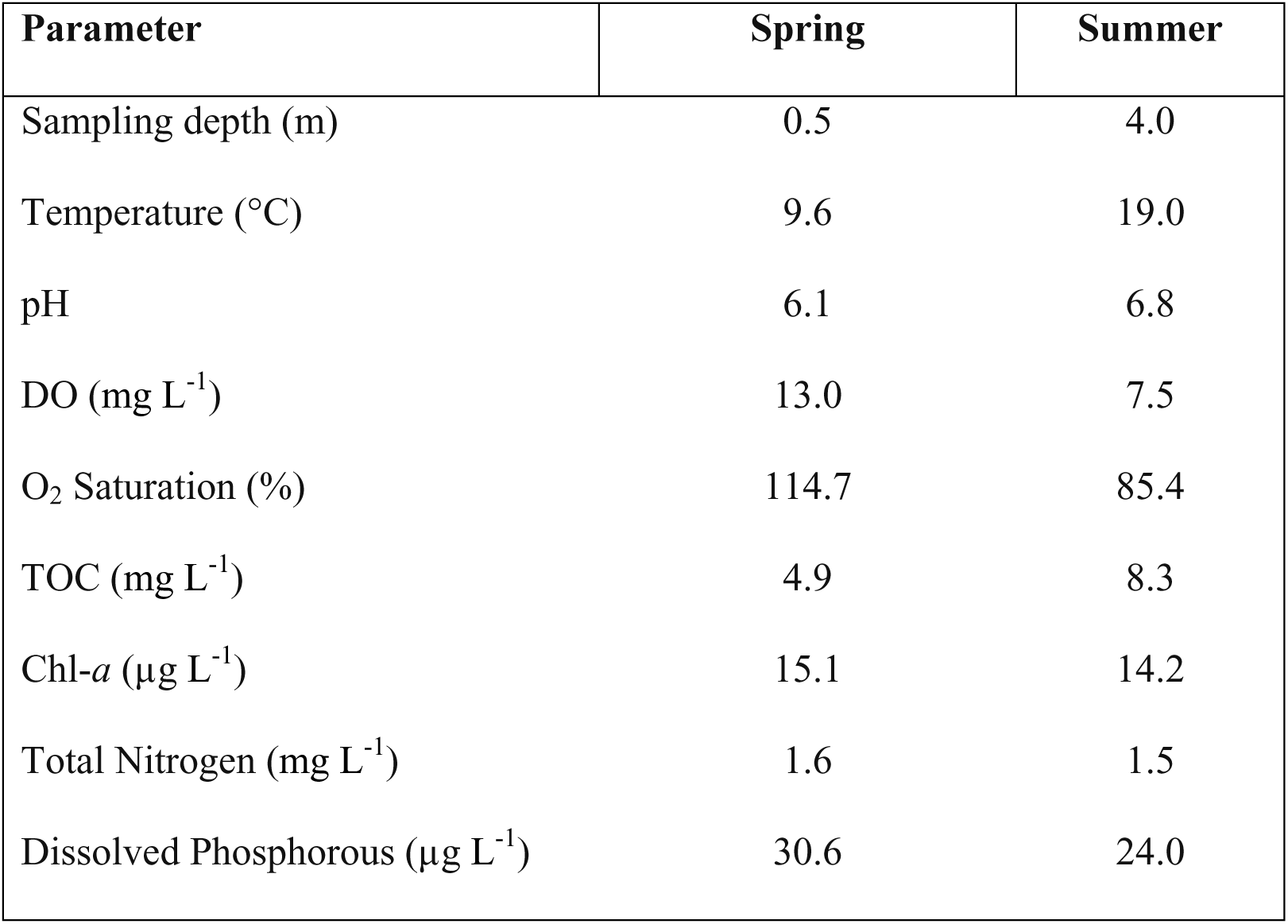
Physical and chemical parameters at the sampling site during spring and summer.

**Table 2.**
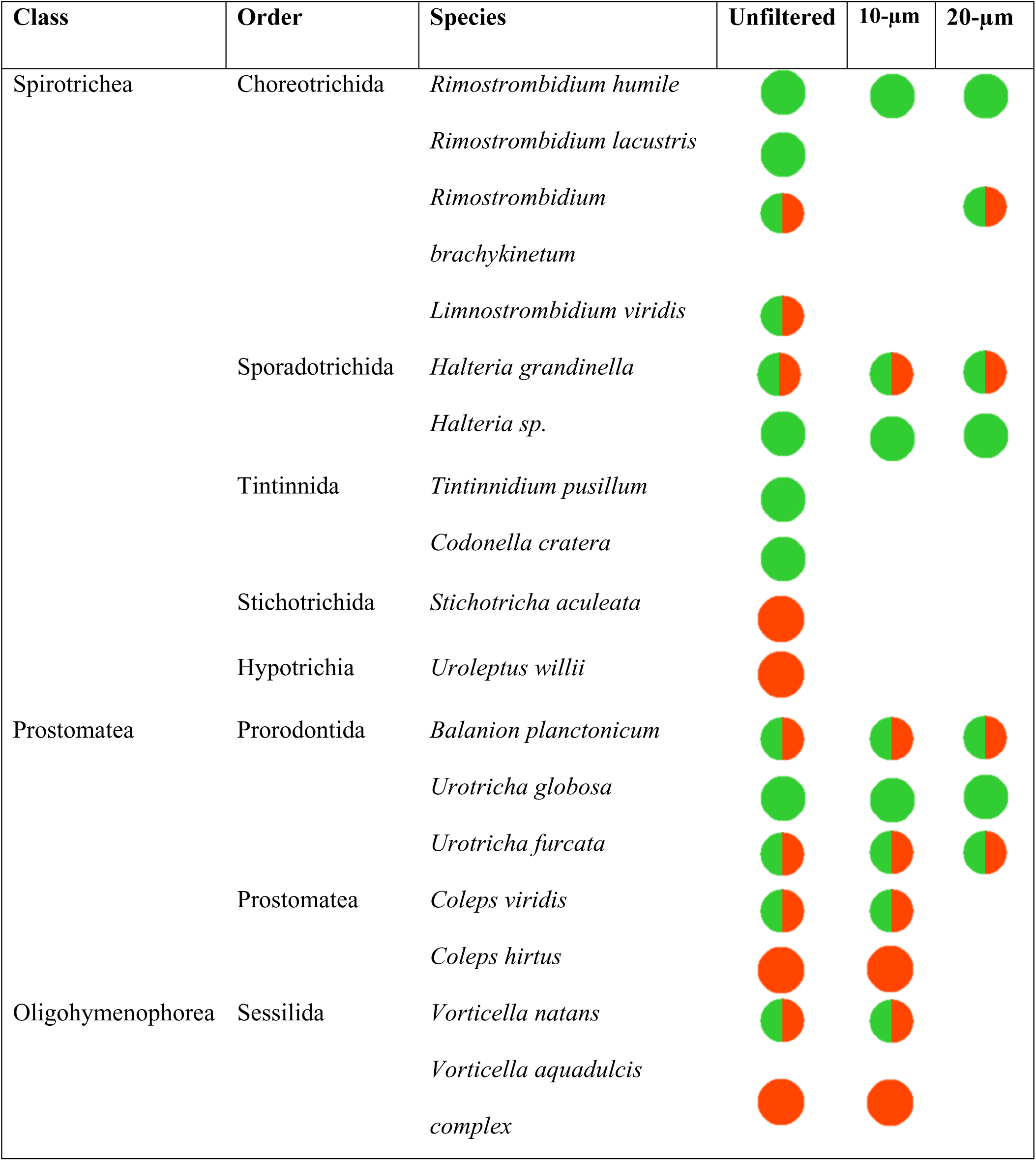
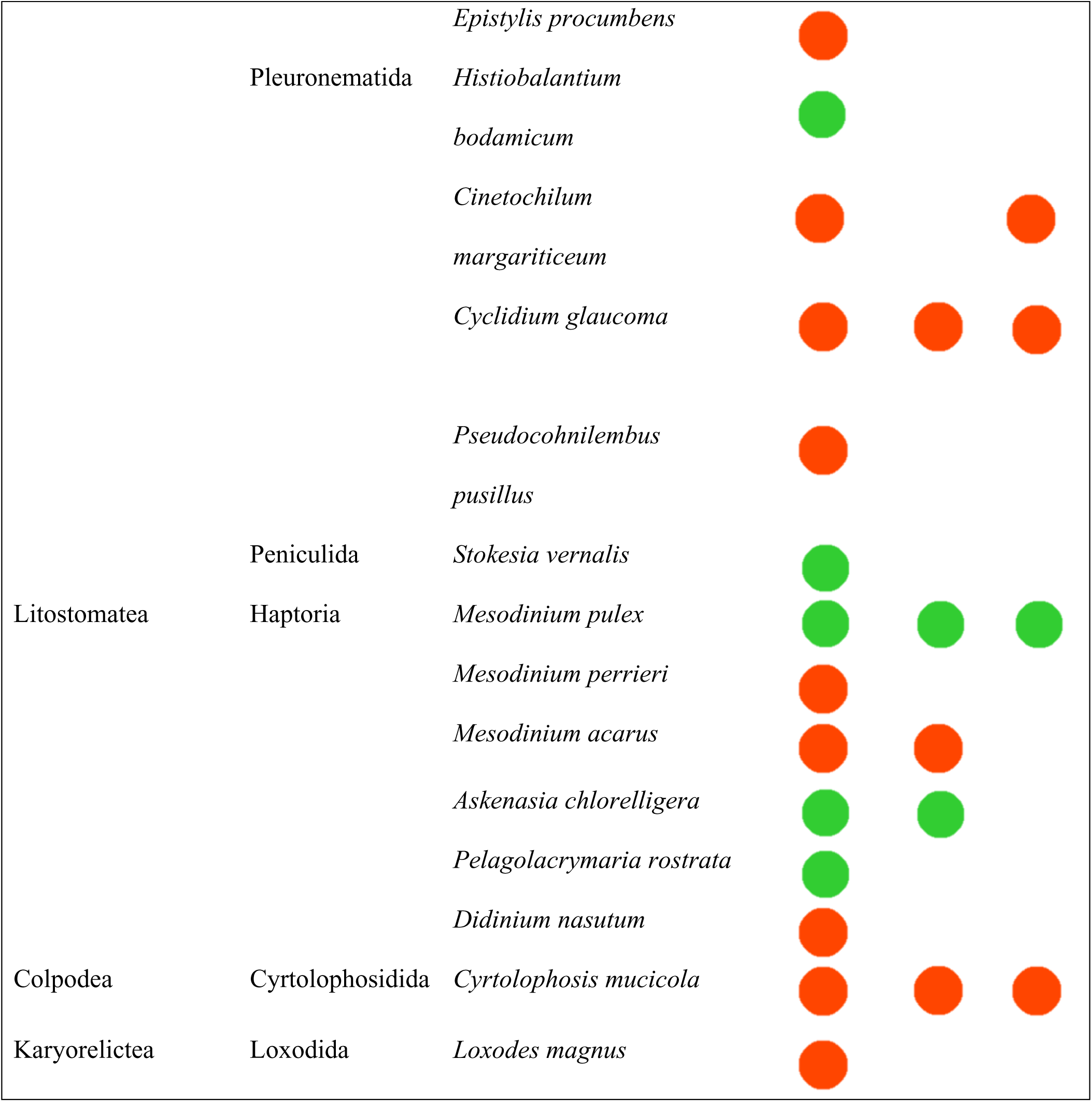
Ciliate taxa detected by QPS during food web manipulation experiments. In different seasons and size fractions. Green for spring, red for summer, green/red for both seasons.

### Response of the spring ciliate community to experimental manipulations: results from microscopic analyses

The size fractionation approach reduced the initial ciliate abundance from the unfiltered (15.1 cells ml^-1^) to the 20-µm (12.5 cells ml ^1^) and 10-µm (9.3 cells ml ^1^) treatments and affected the ciliate community composition (Figures 2A-F). At T0 h, in the unfiltered treatment, *V. natans*, *C. cratera* and *Urotricha* spp. were most abundant, while, in the 20-µm treatment *Urotricha* spp. and *V. natans*, and in the 10-µm treatment *Urotricha* spp., *H. grandinella* and *B. planctonicum* prevailed.

**Figure 2.**
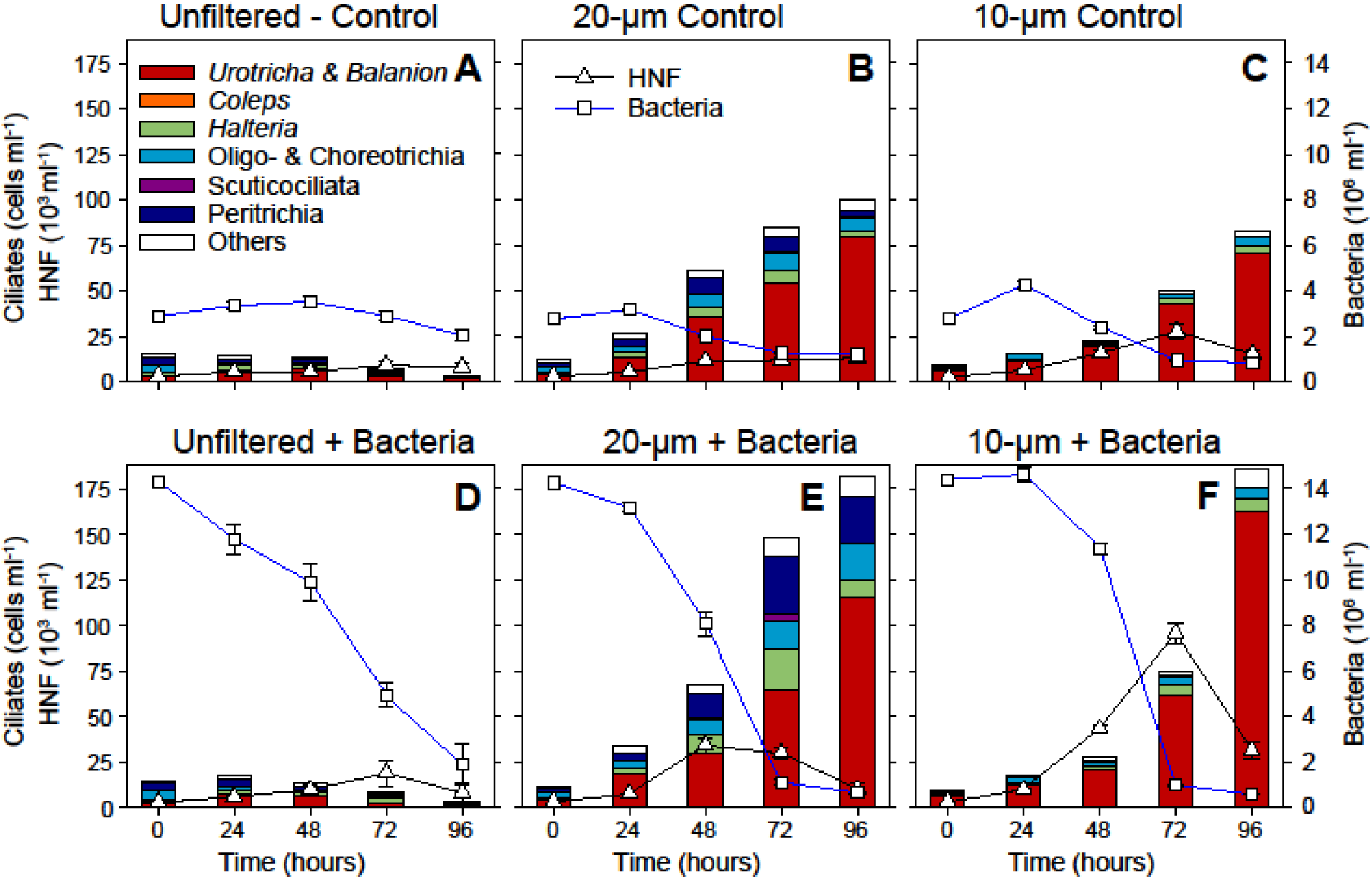
The spring experiment: temporal changes in the abundance of bacteria, HNF and ciliates, and the compositional shifts in the ciliate community (by QPS) between 0 and 96 hours. Error-bars represent the mean values of abundance of different ciliate taxa. Upper panels show the controls (A–C), lower ones the bacterial amended treatments (D–F), from left to right the unfiltered (A, D), the 20-µm (B, E) and the 10-µm treatments (C, F) are shown.

Time-course data from the unfiltered treatment indicated that ciliate communities nearly collapsed both in the control and bacteria-amended treatments at the end of experiment (Figures 2A, D). However, we observed the marked growth response (19 × 10^3^ cells ml^-1^) of HNF at T72 h in the bacteria-amended unfiltered treatment (Figure 2D). In the 20-µm and 10-µm control treatments, a trend in increasing ciliate abundance was observed with 100.9 cells ml^-1^ and 83.2 cells ml^-1^, respectively (Figures 2B, C). The prostomatid flagellate grazers *Urotricha* spp. and *B. planctonicum*, dominated the treatments.

Notably, in bacteria-amended treatments HNF reached their maxima (34.1 × 10^3^ cells ml^-1^) at T48 h in the 20-µm treatment (Figure 2E), and at T72 h in the 10-µm treatment (85.8 × 10^3^ cells ml^-1^, Figure 2F). However, the bacteria were rapidly decimated from 13.2 × 10^6^ cells ml^-1^ to only 1.1 × 10^6^ cells ml^-1^ in 20-µm treatment (Figure 2E). This likely reflected not only the HNF bacterivory, but also the possible contribution of *V. natans* which peaked at T72 h (Figure 2E).

In both the bacteria-amended 10-µm and 20-µm treatments, HNF peaks were followed by conspicuous peaks in ciliate abundance with 186.5 cells ml^-1^ and 182.5 cells ml^-1^, respectively at T96 h. This sharp increase was mainly due to the rapid growth of raptorial flagellate hunters, *B. planctonicum* (10-µm: 81.9 cells ml^-1^, 20-µm: 45.3 cells ml^-1^) and *Urotricha* spp. (10-µm: 80.9 cell ml^-1^, 20-µm: 70.3 cells ml^-1^) (Figures 2E, F). However, in the 20-µm treatment the ciliate assemblage was more diverse, along with remarkable growth of bacterivorous peritrichs and choreotrichs ciliates (Figure 2E). Similarly results of the ANOVA also revealed that time (F=77.623, p<0.001), size fractionation (F=30.514, p<0.001) and prey amendment (F=9.083, p=0.007) had significant effects on ciliate abundance during spring.

### Response of summer ciliate community to experimental manipulation: results from microscopic analyses

During summer, the ciliate community was largely dominated by small ciliates. Therefore, the size fractionation yielded similar initial ciliate abundances in the unfiltered (28.3 cells ml^-1^) and the 20-µm (32.7 cells ml^-1^) treatments (Figures 3A, B), where both the treatments were dominated by *Urotricha* spp., *C. glaucoma*, *Coleps* spp., and *C. margaritaceum* at T0 h (Figures 3A, B). However, in 10-µm ciliate abundance decreased considerably at T0 h (19.4 cells ml^-1^, Figure 3C), being accompanied also by shifts in the composition of ciliates towards tiny species, such as dominant *C. glaucoma,* followed by *C. mucicola* and *Urotricha* spp. (Figure 3C).

**Figure 3.**
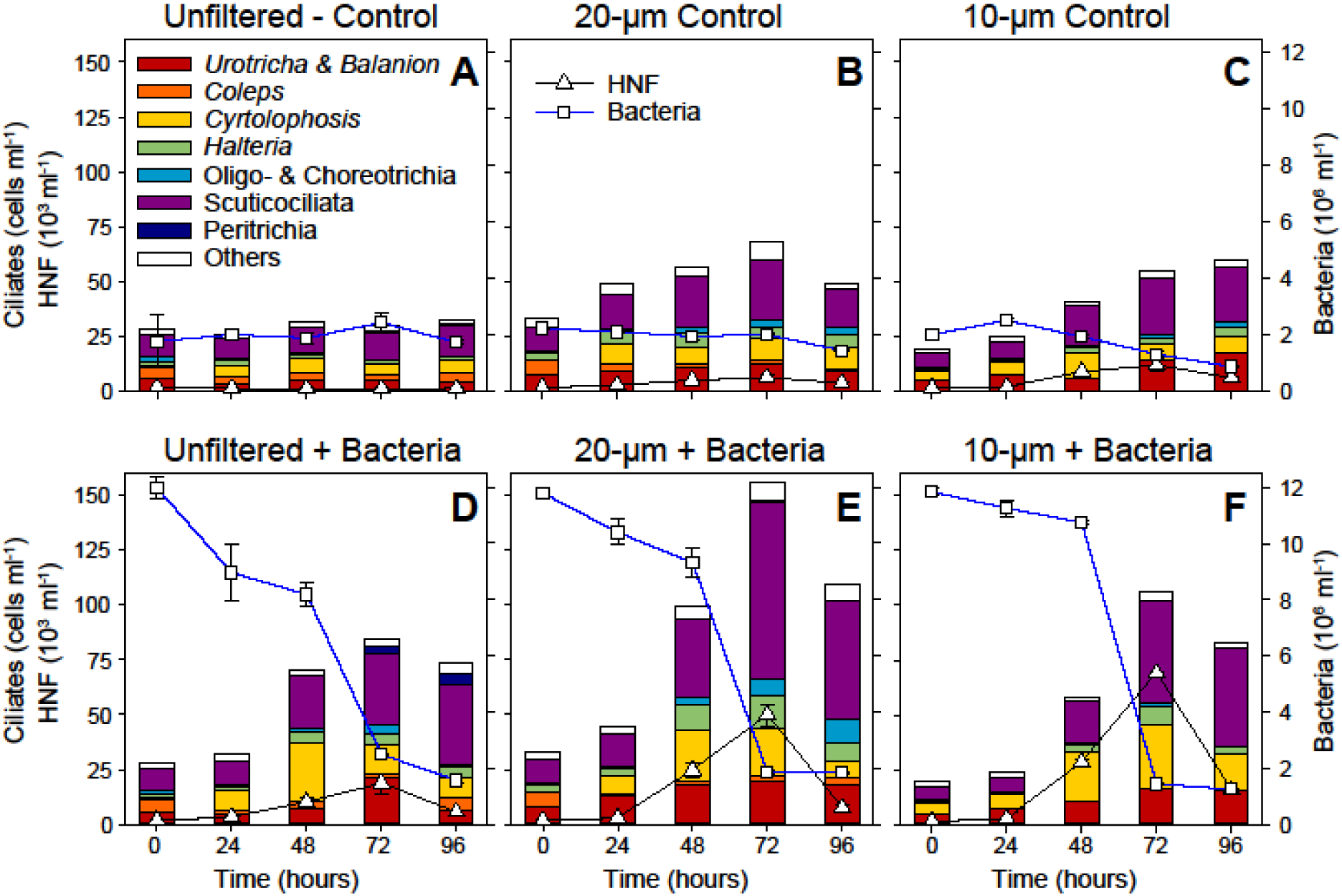
The summer experiment: Temporal changes in the abundance of bacteria, HNF and ciliates, and the compositional shifts in the ciliate community (by QPS) between 0 and 96 hours. Error-bars represent the mean values of abundance of different ciliate taxa. Upper panels show the controls (A–C), lower ones the bacterial amended treatments (D–F), from left to right the unfiltered (A, D), the 20-µm (B, E) and the 10-µm treatments (C, F) are shown.

In the unfiltered treatments, the ciliate community remained relatively stable with a slight increase in ciliate numbers (Figure 3A). In bacterial amended treatment, the ciliate reached 81.3 cells ml^-1^ at T72 h, mainly due to increase of *C. margaritaceum* (19.5 cells ml^-1^) and *Urotricha* spp. (20.1 cells ml^-1^) (Figure 3D). Notably, in the summer prey-amended treatments, maxima of both HNF and ciliate abundance were synchronized at T72 h in 10-µm and 20-µm treatments with 104.3 cells ml^-1^ and 156 cells ml^-1^, respectively, just after the sharp bacterial abundance decline (Figures 3E, F). The bacterivorous scuticociliates *C. glaucoma* (10-µm: 40 cells ml^-1^, 20- µm: 48.4 cells ml^-1^) and *C. margaritaceum* (only in 20-µm: 32.7 cells ml^-1^) and the detritivorous *C. mucicola* (10-µm: 27.4 cells ml^-1^, 20-µm: 23.4 cells ml^-1^) were the prominent species (see Figures 3E, F). Notably control treatments (10-µm: 54.8 cells ml^-1^, 20-µm: 68 cells ml^-1^ at T72h) also showed moderate growth responses (Figures 3B, C). Results of the ANOVA revealed that time (F=33.125, p<0.001), prey amendment (F=18.288, p<0.001) and size fractionation (F=6.827, p=0.006) yielded significant effects on ciliate abundance.

### Shift in the composition of ciliate communities: insights from sequence analysis

In total, we obtained 56,846 high-quality reads affiliated to the phylum Ciliophora comprising 465 distinct ASVs. Spring samples contained 326 ASVs classified in 10 classes, 15 orders, 26 families and 32 genera. No sequences from the bacteria-amended 10-µm treatment at 48 h passed the filtering steps and this sample was not considered for analysis. The dominant groups in the spring ASVs were Hypotrichia (26.1%), Choreotrichia (23.5%), *Askenasia* (16.1%) and *Mesodinium* (9.0%) from the subclass Haptoria (Figure 4). The unfiltered samples showed high proportions of Suctoria at the starting point, which was later replaced by Hypotrichia, especially in the bacteria-amended samples (Figure 4A). On the other hand, 10-µm samples showed high relative abundance of *Mesodinium* at T0 h (40.1%), replaced by *Askenasia* along the experimental time in both treatments (Figure 4C). However, both species were microscopically rare. While *Urotricha spp.* and *B. planctonicum* were the dominant morphotypes, accounting together for more than 85% of total ciliates observed by QPS (Figure 3F) but contributed disproportionally less to the sequencing data. Only *Urotricha* accounted for 3.3% of all ASVs, while *B. planctonicum* together with *P. rostrata*, *C. cratera*, *C. margaritaceum* remained undetected by sequence analysis among the 17 genera identified microscopically. *Mesodinium* also showed higher diversity through sequence analysis (*M. pulex*, *M. rubrum*, *Mesodinium sp*.). In 20-µm samples, Choreotrichia was the dominant group at T96 h (40.6% in control and 51.1% in bacteria-amended samples, Figure 4B).

**Figure 4.**
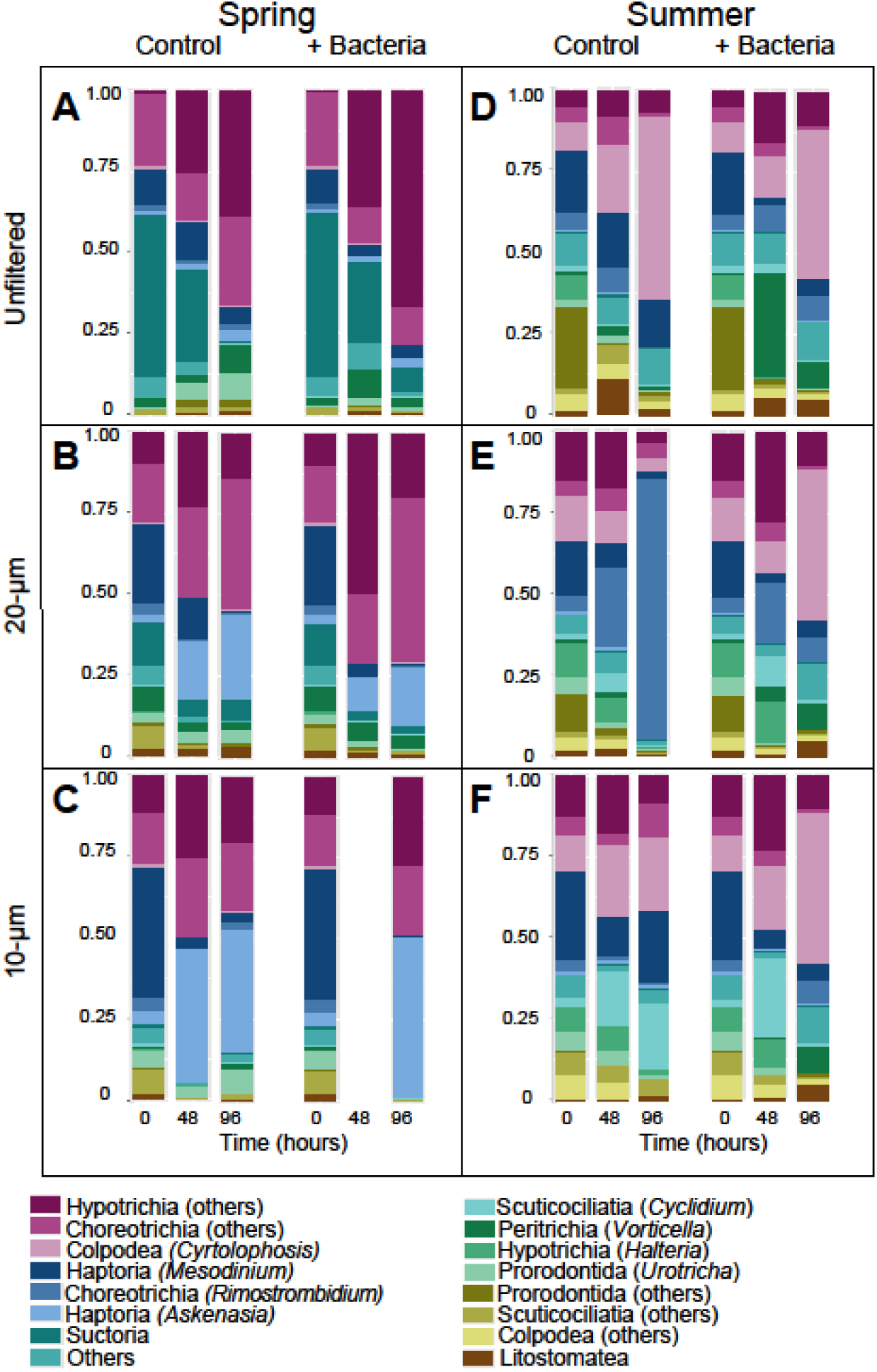
Relative abundance of ciliates in the spring and summer experiments over time in the different size fractions by long-read amplicon sequencing. The results are shown for the respective size fractions in separated columns representing control treatments (Control) and prey-amended treatments (Bacteria +). To align the sequencing results with microscopic counts, ASVs abundance data of microscopically abundant species are presented for respective groups.

During summer, we detected 345 ASVs assigned to 9 classes, 17 orders, 33 families and 34 genera. Choreotrichia (27.5%), Colpodea (19.8%) and Hypotrichia (13.4%) showed the higher relative abundance (Figure 4). *Cyclidium, Cyrtolophosis* and *C. margaritaceum* showed relatively high abundances in summer, and they also dominated the microscopic counts (Figures 3E, F). However, *C. margaritaceum* was undetected in the sequence analysis.

In contrast to spring, Prorodontida (25.5%) was the dominant group at the starting point in the unfiltered samples (Figure 4A), while *Mesodinium* dominated the 10-µm and 20-µm samples (Figures 4B, C). Later it was replaced by *Cyrtolophosis* (>45%) in all bacteria-amended treatments (unfiltered, 10-µm and 20-µm) at T96 h. In bacteria-amended, unfiltered treatment *Vorticella* (32.1%), and in 10-µm, *Cyclidium* showed abundance peak at T48 h (Figure 4A, C). While five distinct ASVs of *Vorticella* (*V. aequilate*, *V. convallaria*, *V. micostoma*, *V. aquadulcis*, *V. astyliformis*) were detected, however, only two species (*V. natans* and *V. aquadulcis* complex) were detected microscopically. Notably, only the 20-µm treatment from the control group showed a different trend with a remarkable increase in the proportion of *Rimostrombidium* (81%), which became the dominant group at the experimental end (Figure 4B). However, microscopically this genus contributed only to 7% of total ciliate abundance (Figure 3E).

dbRDA coupled with PERMANOVA identified seasonality as the main explainer of the variance of the samples (35.51% explained, p = 0.001) by sequencing, as can also be seen from the PCoA multivariate analysis (Figure S2). In order to cluster the samples hierarchically to identify the main taxonomic markers for the factor groups, both seasons showed the best result in the heatmap clustering, clearly highlighting the main taxons responsible for the dissimilarity of the samples between the seasons based on the relative abundance. More taxons stood out in spring than in summer in this cluster analysis, in particular the ones unidentified at genus level (Figure 5).

**Figure 5.**
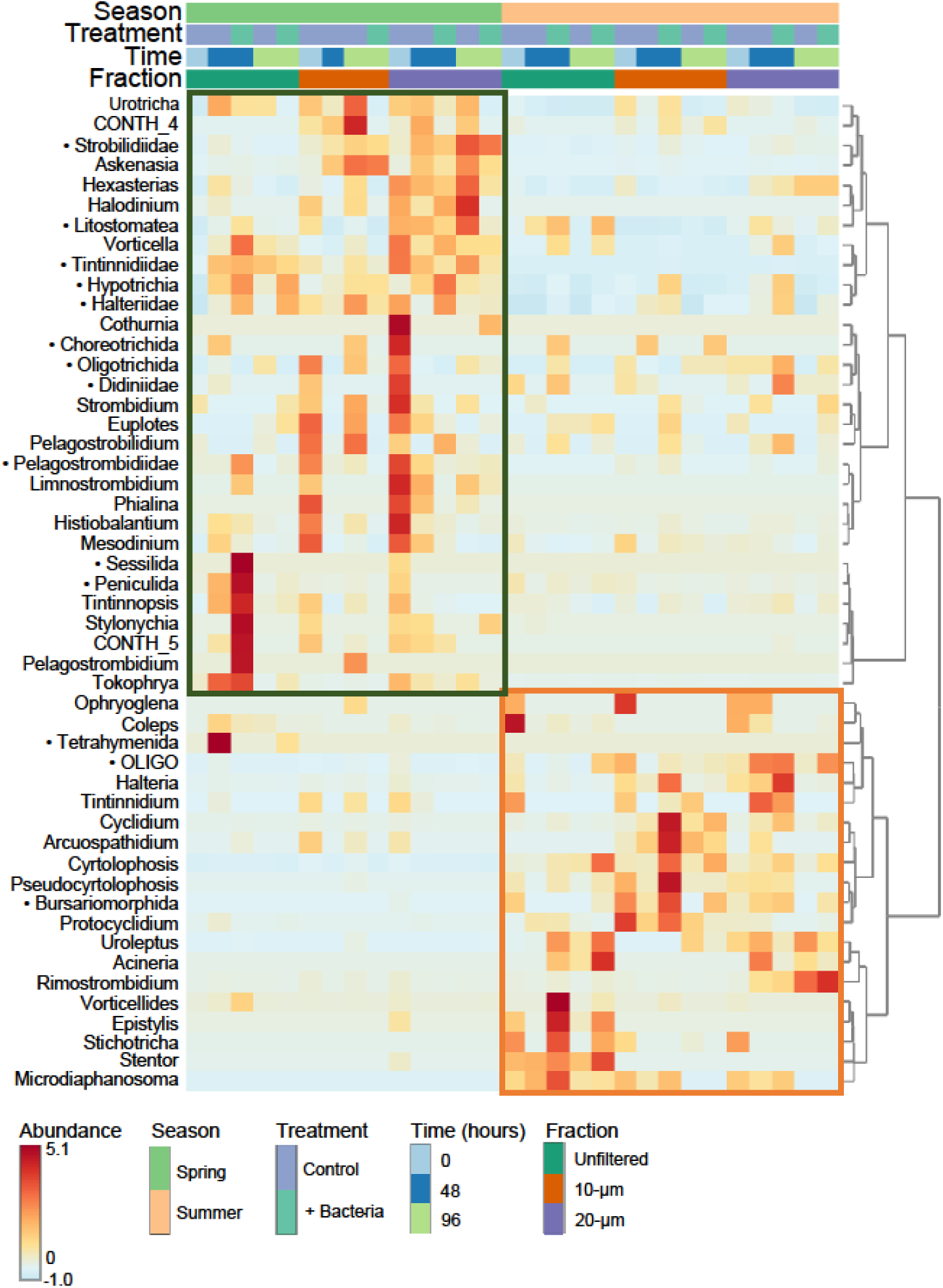
A clustered heatmap showing the variation of relative abundance of the main taxa which distinguishes the samples from different seasons (spring and summer) by long-read amplicon sequencing. Black dots next to the taxa names indicate the unidentified ASVs at genus level.

## DISCUSSION

Two initial seasonally different ciliate communities, reflecting season-specific characteristics of food availability, were experimentally manipulated in the same way (Figure 1) but showed distinct season-specific responses in terms of ciliate abundance and community dynamics.

### Comparison of *in situ* and food web manipulation study during spring

The beginning of spring bloom is usually initiated by the proliferation of small prostome *B. planctonicum*, which peaks immediately at the initiation of phytoplankton bloom but then declines, being replaced by larger ciliates with different life strategies (16, 26). In our study, we collected samples during the mid-phase of the spring bloom, which already showed low numbers of *B. planctonicum* (1 cell ml^-1^). In many lakes (e.g. Lakes Constance and Zurich), *B. planctonicum* is replaced by algivorous/predatory *Urotricha* spp., *H. bodamicum* and *Mesodinium* sp. during the bloom phase (18, 27). However, in our study, bacterial-amended treatments triggered a steep increase of *Urotricha* spp. and *B. planctonicum*. This increase likely reflects the rapidly growing HNF abundance (Figures 2D-F) as a result of bacterial enrichment, dominated by bacterivorous cryptophytes, (28, 29). *B. planctonicum* is a typical r-strategist and responds rapidly to enhanced cryptophyte prey (18, 26, 27) and together with *Urotricha* spp. are typical flagellate predator (26, 28). Indeed, the abundance of HNF peaks in spring plankton, followed by the peak of prostomes (16, 18), further confirming our assumption about the complexity of trophic cascade from HNF to small ciliates (30).

Several studies have attributed the mid-spring decline in ciliate communities, prior to the onset of the clear-water phase, to predation by large grazers such as copepods (calanoids and cyclopoids), cladocerans and rotifers (19, 31, 32). Studies observed that ciliate communities are subject to strong top-down control by mesozooplankton, especially copepods, followed by intraguild predation by daphnids and food competition, which eventually leads to the complete decline of ciliate communities (33–35). Similarly, mesozooplankton including cladocerans, copepods and rotifers showed their annual maximum during spring bloom at our sampling site (17, 36).

Notably, we also observed a strong top-down control in our spring experiment, particularly in unfiltered treatments where ciliates were subjected to severe top-down control by large zooplankton, resulting in a dramatic drop to negligible ciliate numbers towards the experimental end (Figure 2A, D), thus confirming general trends described for late clear water phase (11, 37). Surprisingly, the grazing control of zooplankton even overrode the effect of the considerable bacterial amendment and thus did not induce any net ciliate growth as compared to the marked growth response of HNF (Figure 2D). However, in the filtered treatments (20-µm and 10-µm), ciliate communities were released from the grazing pressure and increased rapidly, especially in bacterial amended treatments, with approximately daily doubling, comparable to the rates of typical pelagic environments (18, 28). Also, the size fractionation had a highly significant effect on ciliate abundance and species composition in spring. Thus, we conclude that the strong top-down control during the bloom phase considerably shaped the ciliate community in our treatments (5, 11, 13, 26, 34, 37). The trends nicely link the outcomes of experiments with the seasonal studies.

### Comparison of *in situ* and food web manipulation study during summer

In previous studies conducted in Římov during the summer (24, 38), ciliates consumed ∼20% of total bacterial production. Ciliate communities were dominated by small-sized (<25 µm) species, especially *H. grandinella*, which was later replaced by *Pelagostrobilidium*, *Coleps* sp. and *C. mucicola*, while *Urotricha* spp. and *C. margaritaceum* made stable contributions (24). We observed similar patterns in ciliate communities in our summer experiment (Figure 3A-F). Notably, the omnivorous/bacterivorous *H. grandinella* was the dominant species in *in-situ* studies and an important omnivorous species linking different trophic levels (24, 39, 40), however, we observed only slight increases in our experiments. Most likely the enhanced bacterial prey availability supported rather bacterivorous/detritivorous species with the observed dominance of *C. glaucoma, C. margaritaceum* and *C. mucicola* (Figure 3D-F).

In all bacteria-amended treatments, *C. glaucoma* was a dominant species (Figure 3D-F), underlying that *Cyclidium* is an efficient bacterivore (24), which is even size-selective (41, 42). Moreover, our results clearly reflected the positive cascading effect of bacterial enrichment on the total ciliate abundance. Besides, the common maxima of both HNF and ciliates abundance were recorded at T72 h, just after the sharp decline of bacterial numbers (Figure 3D-F), indicating the direct transfer of energy from bacteria to both these bacterivore groups.

Generally, grazing pressure of larger zooplankton is assumed to be weaker during summer season (11, 17), and thus ciliate communities should be subjected to a rather moderate top-down control of large zooplankton. However, our experimental treatments suggest considerable effects of summer grazer community. For instance, the results from 10- and 20-µm control treatments indicated considerable levels of grazing on protists, as evidenced by the significantly higher ciliate abundance in these treatments (>60 ml^-1^) compared to unfiltered controlled treatment (31.5 cells ml^-1^). The results of ANOVA also confirmed that size fractionation had significant effects on the ciliate abundance and community composition during summer.

Ecological studies on ciliates have been carried out in both *in situ* and in the laboratory (13, 28, 39, 43). *In situ* experiments mimic natural conditions well but encounter difficulties when conducting experiments on complex planktonic systems, resulting in a lack of precision (44, 45). On the other hand, manipulation of natural plankton communities in the laboratory eliminates the effects of various interfering factors. However, it offers the possibility to accurately measure impacts of specific factors (46, 47), in our case the effects of food availability and top-down control by grazers. Despite some possible shortcomings of the experimental manipulations (48), our study demonstrates its usefulness for providing valuable information on the trophic cascading and top-down control in the seasonally differently shaped microbial communities (47, 49).

### High throughput sequencing based results reflect higher ciliate diversity

Sequencing-based results generally provide a more comprehensive picture of ciliate diversity (21, 50–52). In a study on the composition of ciliates in the Krka River 214 ciliate genera were detected by molecular studies, while only 26 genera were identified microscopically (53).

Similarly, only 30 ciliate genera were observed microscopically and 47 were detected by molecular studies in a mountain lake (54). In line with the previous studies our sequencing data also revealed a much broader ciliate diversity with specific sequences detected in both experiments (Figure 5). However, only 32 ASVs in spring and 36 in summer could be assigned to genus level, which indicates a huge hidden diversity in ciliates.

Numerous studies have also reported large discrepancies between microscopic and molecular abundance data (21, 55, 56). The possession of extremely high copy number, taxonomically uneven distribution and sequence polymorphism of the 18S rRNA genes are among the major reasons for the discrepancy (57, 58). Due to a very high copy number of their 18S rRNA gene ciliates usually dominate the molecular analysis of environmental samples, but their high-read abundance is usually not supported by morphological studies (53, 59, 60). This highlights the obstacle of using sequence-based approaches to determine the abundance of ciliates (56, 58).

Our study confirmed the incongruencies in relative abundance of ciliates between morphological and sequence-based approaches. During spring, Hypotrichia, Choreotrichia, Haptoria (*Aksenasia* and *Mesodinium*) and during summer Choreotrichia, Colpodea and Hypotrichia dominated the sequencing results (Figure 4), but contribute far less to microscopic counts (54, 56, 59), except for Colpodea, dominating morphotype during summer (Figure 3D, F). An obvious reason for this is the extremely high copy number of rRNA genes in ciliates, with the real extremes reported for peritrich ciliates (*Vorticella*) (61). Not surprisingly, *Vorticella* accounted for <1% of the total number of ciliates in bacteria-amended unfiltered treatment at T48 h in summer but accounted for more than 32% of all sequences (Figure 4). Similarly, Choreotrichia (*Rimostrombidium*) also have high gene copy numbers (57, 61). While they represented <10% of cell counts, they accounted for >80% of all sequences in 20-µm control treatment at T96 h during summer.

Interestingly, we found that our molecular data of Scuticociliatia (*Cyclidium*) and Colpodea (*C. mucicola*) projected similarities with the microscopic counts, which corroborated previous studies (21, 62). Our results suggest that sequencing-based analysis could project the abundance of ciliates with lower copy numbers of rRNA genes but are unreliable for ciliates with high copy numbers. Therefore, for ecological studies in ciliates all sequence-based results should be accompanied by a morphological analysis such as QPS.

While sequence-based analysis provides a comprehensive overview of community diversity (55, 56, 63, 64), several challenges remain. These include incomplete and error-prone reference databases (53, 54) and lack of ciliate sequences from cultured representatives (21, 54).

Consequently, our results were also influenced by these factors. Seven species were undetected in our molecular analysis, specifically, *B. planctonicum* in spring and *C. margaritaceum* in summer, although they dominated in microscopic counts. This discrepancy is probably due to primer bias or the presence of cryptic species complexes for which there are no representative sequences in public databases (21). Thus, we conclude that the combination of novel sequencing approaches along with classical QPS and fluorescence microscopy are still mandatory. The combination of these techniques provides deeper and more precise insights into ecological aspects, such as community dynamics, rapid changes in abundance of ciliate species and their food preferences under different experimental regimes and seasonal settings.

## MATERIALS AND METHODS

### Study area and sampling site

Samples were collected from a deep, meso-eutrophic Římov Reservoir (48° 51N, 14°29E), located south of České Budějovice, Czech Republic. The reservoir is dimictic with a total surface area of 210 ha and the maximum and mean depth of 42 m and 16 m, respectively (65, 66).

Samples for two experiments were taken in the spring (19^th^ April) and summer (22^nd^ August) of the year 2022 from the epilimnion of the dam area, representing the deepest part of the reservoir. Vertical profiles of temperature, pH, and dissolved oxygen (DO) concentration were measured at the sampling sites by a YSI EXO II multiparameter probe (Yellow Springs Instruments, USA).

### Sampling and experimental manipulation

Forty liters of water were taken from the epilimnion (0.5 m depth) in spring and from a slightly deeper layer (4 m depth) in summer, due to the low DO concentration in the surface layer covered by cyanobacteria. The samples were collected by Friedinger sampler (Šramhauser spol. s.r.o., Czech Republic). Two liters of an untreated subsample were separated for further analysis of chemical parameters including chlorophyll-*a* (Chl-*a*), total organic carbon (TOC), total nitrogen and dissolved phosphorus (DP) (67).

To test for trophic cascading effects of different grazers, we manipulated top-down controlling factors using the size-fractionation approach similar to that described elsewhere (28). Size fractionation was done by gravity filtration through 20-µm and 10-µm pore-size filters (diameter 142 mm, Sterlitech, USA) and yielded treatments of different food web complexity (Figure 1): (i) the 10-µm fraction contained mainly prokaryotes, small flagellates (< 10-μm) and nano-sized ciliates (< 15-μm); (ii) a more complex 20-µm fraction, which contained additionally also micro-sized ciliates (≥15–30-μm) and large-sized flagellates (10–20-μm), and (iii) compared to an unfiltered whole water sample serving as control with complex natural plankton community. This experimental setup allowed releasing the microbial communities from grazing pressure by larger zooplankton or larger protists, thus gradually representing more and more simplified microbial food webs; from unfiltered to the 20-µm and 10-µm filtered treatments. The water from all the size-fractions was collected in 2-L bottles, and a total of 6 bottles (12 liters) were prepared for each treatment (Figure 1).

In addition, to manipulate bottom-up factors aimed at accelerating trophic cascading within the microbial food webs, one half of all the types of triplicate treatments were amended by pre-cultured bacterial prey (a mixture of isolated strains *Limnohabitans planktonicus* and *L. parvus*) isolated from the reservoir. These bacteria have been proven to be suitable food resources busting the growth of natural communities of bacterivorous protists (28, 68). The amount of the added bacteria was ca. 5-6 times more than the natural background bacterial abundance (yielding the final concentration of 15–20 × 10^6^ cells ml^-1^). All treatments were incubated in the dark at 16° C to set a comparable growth potential of ciliates in both experiments conducted over a period of 96 hours and subsamples of defined volume were aseptically taken every 24 hours at a clean bench.

### Enumeration of HNF and bacteria

Fifteen to twenty ml sub-samples were daily collected from each triplicate and fixed with formaldehyde (2% final concentration). The total bacterial cell counts were quantified via flow cytometry in samples stained with SYBR Green using a CytoFLEX S flow cytometer (Beckman Coulter), equipped with a blue laser and bandpass filters 525/40 and 690/50. For the enumeration of HNF, 5−10 ml subsamples were stained with DAPI (4′,6-diamidino-2-phenylindole, final concentration of 1 μg ml^−1^) and filtered onto 1-μm pore-sized black polycarbonate membrane filters (Sterlitech, USA) and total HNF were counted under an epifluorescence microscope at a magnification of 1000× according to (28).

### Enumeration and identification of ciliates

Samples (100 ml) were daily collected from all triplicate treatments and divided into two 50 ml subsamples. One 50-ml subsample was fixed with formaldehyde (2% final concentration), stained with DAPI and filtered onto black 1-μm pore-sized polycarbonate membrane filters (Sterlitech, USA) for total ciliate counts and identification via epifluorescence microscopy (5, 28, 39, 42). For taxonomic resolution of the morphotypes, we used quantitative protargol staining method (QPS) with second 50-ml subsample that was fixed with Bouin’s solution (5% final concentration) and filtered onto 0.8-μm pore-size nitrocellulose filters (Sartorius, Germany) and stained following the protocol of Skibbe (69), with modifications suggested elsewhere (70). Filters were inspected under bright-field microscopy (Olympus BX50; Japan). Cell was taxonomically identified by using the taxonomic keys of Foissner et al., (71–73) and Foissner and Berger (74)

### DNA extraction, PCR amplification and PacBio sequencing

Three subsamples were collected through the course of the experiment, at times 0 h, 48 h and 96 h. A 350-ml subsample was taken from each replicate and the triplicates from the individual treatments were pooled, resulting in a total volume of 1,050 ml per treatment. Biomass was collected on 0.22-μm polyether sulfone membrane filters (MILLIPORE EXPRESS, Germany) and stored with 1 ml of DNA/RNA Shield at -80°C. For long-read sequencing, DNA was extracted with the Quick-DNA High Molecular Weight MagBead Kit (Zymo Research, USA) according to the manufacturer’s instructions and the concentration was quantified using a Qubit fluorometer (Invitrogen, USA). Extracted DNA was sent to Rush University Genomics and Microbiome Core Facility (Chicago, USA) and there the amplification was conducted with the primer pairs EukV4F (55) and Euk21R (75) to amplify the 18S rRNA, ITS1, ITS2 and a part of the 28S gene following the previously established protocol Latz (76). An equimolar library was prepared using Fluidigm barcodes, followed by PacBio library preparation with the SMRTbell system for high-fidelity long-read sequencing. The sequencing was performed on a Sequel II platform using a SMRT 8M cell with a 30-hour movie runtime. The resulting sequences were demultiplexed and dereplicated at the Rush University Research Bioinformatics Core Facility using DADA2 script specifically designed for PacBio long-amplicon data (77). Chimeric sequences were identified and filtered using the BimeraDenovo algorithm, with a minimum fold parent-over-abundance threshold set to 3.5. Generated long-read sequences were assessed to further quality filtering, primer removal, trimming and denoising using the DADA2 R package, largely following the long-read workflow previously established for the same approach (76, 77). The 18S rRNA gene database PR2 version 5.0.1 (78), was used as training set for taxonomic classification with assignTaxonomy of DADA2. Sequencing data generated in this study are available at the European Nucleotide Archive (ENA) under the BioProject number PRJEB86180.

### Statistical analysis

An ANOVA was performed to examine the effects of time, size fraction and prey amendment (explanatory variables) on ciliate abundance. For long-read amplicon sequences, all amplicon sequence variant (ASV) belonging to the group Ciliophora were considered for further analysis in RStudio (79) and MicrobiomeAnalyst (80). Diversity indices were evaluated using “vegan” package (81) and a two-way Analysis of Variance (ANOVA) for variation in the Shannon diversity index with factors being treatments (control or bacterial amended), size fraction (unfiltered, 10-µm, 20-µm), and different time points (0h, 48h, 96h). We performed the post-hoc Tukey’s honestly significant difference (HSD) test on significant results of the ANOVA to identify which specific groups differed significantly from each other. The β-diversity comparison was performed by principal coordinate analysis (PCoA) using Bray-Curtis dissimilarity and an hierarchical clustering ordination based on the relative abundance of the main genus on the grouping factor were visualized by heatmap using Ward algorithm. Distance-based redundancy analysis (dbRDA) was used to evaluate the dissimilarity between the samples using miaverse package (82) and the permutational ANOVA (PERMANOVA) estimated the statistical significance of the differences in community composition and the variation explained by each grouping factor (treatments, size fraction and time points) using adonis2 function from the vegan package.

## AUTHOR CONTRIBUTIONS

IM, KŠ and KP conceptualization of the study. IM, KŠ, UA, VK, KP sampling, manipulation experiment and total counts. UA, KŠ and BS morphological analysis. IM, UA, CCPD, TS, APS and P-A B DNA extraction and sequence analysis. KŠ and CCPD data visualization. UA wrote the manuscript with input from all the authors. All authors contributed to the critical revisions of the article and approved the submitted version.

## ACKNOWLEDGEMENTS

We are thankful to Dagmara Sirová, Radka Malá, Jakub Psohlavec, Lenka Kosová and Hana Kratochvílová for helping with the experimental set up and excellent laboratory assistance. This work was supported by the research Grant 22-35826 K (Grant Agency of the Czech Republic) awarded to IM. The funding bodies had no role in study design, data collection and analysis, interpretation of data and preparation of the manuscript.

## REFERENCES

1. Pierce RW, Turner JT. Ecology of planktonic ciliates in marine food webs. Reviews in Aquatic Sciences. 1992;6(2):139–81.

2. Gao F, Warren A, Zhang Q, Gong J, Miao M, Sun P, et al. The all-data-based evolutionary hypothesis of ciliated protists with a revised classification of the phylum Ciliophora (Eukaryota, Alveolata). Scientific Reports. 2016;6(1):1–14.

3. Azam F, Ammerman JW. Cycling of organic matter by bacterioplankton in pelagic marine ecosystems: microenvironmental considerations. In: Fasham MJR, editor. Flows of energy materials in marine ecosystems: Theory practice. 13: Springer; 1984. p. 345-60.

4. Weisse T, Müller H, Pinto-Coelho RM, Schweizer A, Springmann D, Baldringer G. Response of the microbial loop to the phytoplankton spring bloom in a large prealpine lake. Limnology Oceanography. 1990;35(4):781–94.

5. Zingel P, Agasild H, Noges T, Kisand V. Ciliates are the dominant grazers on pico-and nanoplankton in a shallow, naturally highly eutrophic lake. Microbial Ecology. 2007;53:134–42.

6. Muylaert K, Van der Gucht K, Vloemans N, Meester LD, Gillis M, Vyverman W. Relationship between bacterial community composition and bottom-up versus top-down variables in four eutrophic shallow lakes. Applied Environmental Microbiology. 2002;68(10):4740–50.

7. Zingel P, Agasild H, Karus K, Kangro K, Tammert H, Tõnno I, et al. The influence of zooplankton enrichment on the microbial loop in a shallow, eutrophic lake. European Journal of Protistology. 2016;52:22–35.

8. Sherr E, Sherr B. Bacterivory and herbivory: key roles of phagotrophic protists in pelagic food webs. Microbial Ecology. 1994;28:223–35.

9. Lu X, Weisse T. Top-down control of planktonic ciliates by microcrustacean predators is stronger in lakes than in the ocean. Scientific Reports. 2022;12(1):10501.

10. Müller H, Schöne A, Pinto-Coelho R, Schweizer A, Weisse T. Seasonal succession of ciliates in Lake Constance. Microbial Ecology. 1991;21:119–38.

11. Sommer U, Adrian R, De Senerpont Domis L, Elser JJ, Gaedke U, Ibelings B, et al. Beyond the plankton ecology group(peg) model: mechanisms driving plankton succession. Annual Review of Ecology, Evolution Systematics. 2012;43(429):2012.

12. Biyu S. A comparative study on planktonic ciliates in two shallow mesotrophic lakes (China): species composition, distribution and quantitative importance. Hydrobiologia. 2000;427:143–53.

13. Li J, Chen F, Liu Z, Zhao X, Yang K, Lu W, et al. Bottom-up versus top-down effects on ciliate community composition in four eutrophic lakes (China). European Journal of Protistology. 2016;53:20–30.

14. Pfister G, Auer B,Arndt H.i Pelagic ciliates (Protozoa, Ciliophora) of different brackish and freshwater lakes—a community analysis at the species level. Limnologica. 2002;32(2):147–68.

15. Beaver JR, Crisman TL. The role of ciliated protozoa in pelagic freshwater ecosystems. Microbial Ecology. 1989;17:111–36.

16. Šimek K, Nedoma J, Znachor P, Kasalický V, Jezbera J, Hornňák K, et al. A finely tuned symphony of factors modulates the microbial food web of a freshwater reservoir in spring. Limnology and Oceanography. 2014;59(5):1477–92.

17. Park H, Shabarova T, Salcher MM, Kosová L, Rychtecký P, Mukherjee I, et al. In the right place, at the right time: the integration of bacteria into the Plankton Ecology Group model. Microbiome. 2023;11(1):112.

18. Posch T, Eugster B, Pomati F, Pernthaler J, Pitsch G, Eckert EM. Network of interactions between ciliates and phytoplankton during spring. Frontiers in Microbiology. 2015;6:1289.

19. Sonntag B, Posch T, Klammer S, Teubner K, Psenner R. Phagotrophic ciliates and flagellates in an oligotrophic, deep, alpine lake: contrasting variability with seasons and depths. Aquatic Microbial Ecology. 2006;43(2):193–207.

20. Zingel P, Nõges T. Seasonal and annual population dynamics of ciliates in a shallow eutrophic lake. Fundamental Applied Limnology. 2010;176(2):133.

21. Pitsch G, Bruni EP, Forster D, Qu Z, Sonntag B, Stoeck T, et al. Seasonality of planktonic freshwater ciliates: are analyses based on V9 regions of the 18S rRNA gene correlated with morphospecies counts? Frontiers in Microbiology. 2019;10:248.

22. Cleven EJ. Pelagic ciliates in a large mesotrophic lake: seasonal succession and taxon-specific bacterivory in lake Constance. International Review of Hydrobiology. 2004;89(3):289–304.

23. Müller H. *Pseudobalanion planctonicum* (Ciliophora, Prostomatida): ecological significance of an algivorous nanociliate in a deep meso-eutrophic lake. Journal of Plankton Research. 1991;13(1):247–62.

24. Šimek K, Bobková J, Macek M, Nedoma J, Psenner R. Ciliategrazing on picoplankton in a eutrophic reservoir during the summer phytoplankton maximum: A study at the species and community level. Limnology and Oceanography. 1995;40(6):1077–90.

25. Jürgens K, Skibbe O, Jeppesen E. Impact of metazooplankton on the composition and population dynamics of planktonic ciliates in a shallow, hypertrophic lake. Aquatic Microbial Ecology. 1999;17(1):61–75.

26. Aberle N, Lengfellner K, Sommer U. Spring bloom succession, grazing impact and herbivore selectivity of ciliate communities in response to winter warming. Oecologia. 2007;150:668–81.

27. Müller H, Schlegel A. Responses of three freshwater planktonic ciliates with different feeding modes to cryptophyte and diatom prey. Aquatic Microbial Ecology. 1999;17(1):49–60.

28. Šimek K, Grujčić V, Mukherjee I, Kasalický V, Nedoma J, Posch T, et al. Cascading effects in freshwater microbial food webs by predatory Cercozoa, Katablepharidacea and ciliates feeding on aplastidic bacterivorous cryptophytes. FEMS Microbiology Ecology. 2020;96(10):fiaa121.

29. Grujčić V, Nuy JK, Salcher MM, Shabarova T, Kasalicky V, Boenigk J, et al. Cryptophyta as major bacterivores in freshwater summer plankton. The ISME Journal. 2018;12(7):1668–81.

30. Piwosz K, Mukherjee I, Salcher MM, Grujčić V, Šimek K. CARD-FISH in the sequencing era: opening a new universe of protistan ecology. Frontiers in Microbiology. 2021;12:640066.

31. Burns CW, Schallenberg M. Calanoid copepods versus cladocerans: consumer effects on protozoa in lakes of different trophic status. Limnology Oceanography. 2001;46(6):1558–65.

32. Jack JD, Gilbert JJ. Effects of metazoan predators on ciliates in freshwater plankton communities 1. Journal of Eukaryotic Microbiology. 1997;44(3):194–9.

33. Tirok K, Gaedke U. Regulation of planktonic ciliate dynamics and functional composition during spring in Lake Constance. Aquatic Microbial Ecology. 2007;49(1):87–100.

34. Zöllner E, Santer B, Boersma M, Hoppe HG, Jürgens K. Cascading predation effects of Daphnia and copepods on microbial food web components. Freshwater Biology. 2003;48(12):2174–93.

35. Jürgens K, Jeppesen E. The impact of metazooplankton on the structure of the microbial food web in a shallow, hypertrophic lake. Journal of Plankton Research. 2000;22(6):1047–70.

36. Mukherjee I, Grujčić V, Salcher MM, Znachor P, Seďa J, Devetter M, et al. Integrating depth-dependent protist dynamics and microbial interactions in spring succession of a freshwater reservoir. Environmental Microbiome. 2024;19(1):31.

37. Kath NJ, Thomas MK, Gaedke U. Mysterious ciliates: seasonally recurrent and yet hard to predict. Journal of Plankton Research. 2022;44(6):891–910.

38. Macek M, Šmek K, Pernthaler J, Vyhnalek V, Psenner R. Growth rates of dominant planktonic ciliates in two freshwater bodies of different trophic degree. Journal of Plankton Research. 1996;18(4):463–81.

39. Šimek K, Grujčić V, Nedoma J, Jezberová J, Šorf M, Matoušů A, et al. Microbial food webs in hypertrophic fishponds: Omnivorous ciliate taxa are major protistan bacterivores. Limnology and Oceanography. 2019;64(5):2295–309.

40. Forster D, Qu Z, Pitsch G, Bruni EP, Kammerlander B, Pröschold T, et al. Lake ecosystem robustness and resilience inferred from a climate-stressed protistan plankton network. Microorganisms. 2021;9(3):549.

41. Posch T, Jezbera J, Vrba J, Šimek K, Pernthaler J, Andreatta S, et al. Size selective feeding in Cyclidium glaucoma (Ciliophora, Scuticociliatida) and its effects on bacterial community structure: a study from a continuous cultivation system. Microbial Ecology. 2001;42:217–27.

42. Šimek K, Macek M, Pernthaler J, Straškrabová V, Psenner R. Can freshwater planktonic ciliates survive on a diet of picoplankton? Journal of Plankton Research. 1996;18(4):597–613.

43. Rychert K, Wielgat-Rychert M, Szczurowska D, Myszka M, Bochyńska M, Krawiec K. The importance of ciliates as a trophic link in shallow, brackish and eutrophic lakes. Polish Journal of Ecology. 2012;60(4):767–76.

44. Vijverberg J. Culture techniques for studies on the growth, development and reproduction of copepods and cladocerans under laboratory and in situ conditions: a review. Freshwater Biology. 1989;21(3):317–73.

45. Agasild H, Zingel P, Karus K, Kangro K, Salujõe J, Noges T. Does metazooplankton regulate the ciliate community in a shallow eutrophic lake? Freshwater Biology. 2013;58(1):183–91.

46. Weisse T, Montagnes DJ. Ecology of planktonic ciliates in a changing world: Concepts, methods, and challenges. Journal of Eukaryotic Microbiology. 2022;69(5):e12879.

47. Montagnes DJ, Barbosa AB, Boenigk J, Davidson K, Jürgens K, Macek M, et al. Selective feeding behaviour of key free-living protists: avenues for continued study. Aquatic Microbial Ecology. 2008;53(1):83–98.

48. Mullen AD, Treibitz T, Roberts PL, Kelly EL, Horwitz R, Smith JE, et al. Underwater microscopy for in situ studies of benthic ecosystems. Nature Communications. 2016;7(1):12093.

49. Weisse T. Functional diversity of aquatic ciliates. European Journal of Protistology. 2017;61:331–58.

50. Zhao Y, Langlois GA. Ciliate morpho-taxonomy and practical considerations before deploying metabarcoding to Ciliate community diversity surveys in urban receiving waters. Microorganisms. 2022;10(12):2512.

51. Abraham JS, Sripoorna S, Maurya S, Makhija S, Gupta R, Toteja R. Techniques and tools for species identification in ciliates: a review. International Journal of Systematic Evolutionary Microbiology. 2019;69(4):877–94.

52. Dong J, Shi F, Li H, Zhang X, Hu X, Gong J. SSU rDNA sequence diversity and seasonally differentiated distribution of nanoplanktonic ciliates in neritic Bohai and Yellow Seas as revealed by T-RFLP. PLoS One. 2014;9(7):e102640.

53. Kulaš A, Gulin V, Kepčija RM, Žutinić P, Perić MS, Orlić S, et al. Ciliates (Alveolata, Ciliophora) as bioindicators of environmental pressure: A karstic river case. Ecological Indicators. 2021;124:107430.

54. Stoeck T, Breiner HW, Filker S, Ostermaier V, Kammerlander B, Sonntag B. A morphogenetic survey on ciliate plankton from a mountain lake pinpoints the necessity of lineage-specific barcode markers in microbial ecology. Environmental Microbiology. 2014;16(2):430–44.

55. Stoeck T, Bass D, Nebel M, Christen R, Jones MD, BREINER HW, et al. Multiple marker parallel tag environmental DNA sequencing reveals a highly complex eukaryotic community in marine anoxic water. Molecular Ecology. 2010;19:21–31.

56. Medinger R, Nolte V, Pandey RV, Jost S, Ottenwälder B, Schlötterer C, et al. Diversity in a hidden world: potential and limitation of next-generation sequencing for surveys of molecular diversity of eukaryotic microorganisms. Molecular Ecology. 2010;19:32–40.

57. Wang Y, Jiang Y, Liu Y, Li Y, Katz LA, Gao F, et al. Comparative studies on the polymorphism and copy number variation of mtSSU rDNA in ciliates (Protista, Ciliophora): implications for phylogenetic, environmental, and ecological research. Microorganisms. 2020;8(3):316.

58. Gong J, Dong J, Liu X, Massana R. Extremely high copy numbers and polymorphisms of the rDNA operon estimated from single cell analysis of oligotrich and peritrich ciliates. Protist. 2013;164(3):369–79.

59. Yu L, Zhang W, Liu L, Yang J. Determining microeukaryotic plankton community around Xiamen Island, southeast China, using Illumina MiSeq and PCR-DGGE techniques. PLoS One. 2015;10(5):e0127721.

60. Dopheide A, Lear G, Stott R, Lewis G. Relative diversity and community structure of ciliates in stream biofilms according to molecular and microscopy methods. Applied Environmental Microbiology. 2009;75(16):5261–72.

61. Gong J, Dong J, Liu X, Massana R. Extremely high copy numbers and polymorphisms of the rDNA operon estimated from single cell analysis of oligotrich and peritrich ciliates. Protist. 2013;164(3):369–79.

62. Gertler C, Näther DJ, Gerdts G, Malpass MC, Golyshin PN. A mesocosm study of the changes in marine flagellate and ciliate communities in a crude oil bioremediation trial. Microbial Ecology. 2010;60:180–91.

63. Liu X, Gong J. Revealing the diversity and quantity of peritrich ciliates in environmental samples using specific primer-based PCR and quantitative PCR. Microbes Environments. 2012;27(4):497–503.

64. Caron DA, Countway PD, Savai P, Gast RJ, Schnetzer A, Moorthi SD, et al. Defining DNA-based operational taxonomic units for microbial-eukaryote ecology. Applied Environmental Microbiology 2009;75(18):5797–808.

65. Blabolil P, Ricard D, Peterka J, Říha M, Jůza T, Vašek M, et al. Predicting asp and pikeperch recruitment in a riverine reservoir. Fisheries Research. 2016;173:45–52.

66. Znachor P, Hejzlar J, Vrba J, Nedoma J, Seda J, Šimek K, et al. Brief history of long-term ecological research into aquatic ecosystems and their catchments in the Czech Republic. Part I: Manmade reservoirs Institute of Hydrobiology, BC CAS, České Budějovice. 2016;1:88.

67. Znachor P, Nedoma J, Hejzlar J, Seďa J, Kopáček J, Boukal D, et al. Multiple long-term trends and trend reversals dominate environmental conditions in a man-made freshwater reservoir. Science of the Total Environment. 2018;624:24–33.

68. Šimek K, Grujčić V, Hahn MW, Horňák K, Jezberová J, Kasalický V, et al. Bacterial prey food characteristics modulate community growth response of freshwater bacterivorous flagellates. Limnology and Oceanography. 2018;63(1):484–502.

69. Skibbe O. An improved quantitative protargol stain for ciliates and other planktonic protists. Archiv für Hydrobiologie. 1994;30:339–47.

70. Pfister G, Sonntag B, Posch T. Comparison of a direct live count and an improved quantitative protargol stain (QPS) in determining abundance and cell volumes of pelagic freshwater protozoa. Aquatic Microbial Ecology. 1999;18(1):95–103.

71. Foissner W, Berger H, Kahoman F. Taxonomische und ökologische revision der ciliaten des saprobiensystems. Band III: Hymenostomata, Prostomatida, Nassulida. . Informationsberichte des Bayerischen Landesamt fur Wasserwirtschaft. 1994.

72. Foissner W, Berger H, Schaumburg J. Identification and ecology of limnetic plankton ciliates. Munich: Landesamtes für Wissenschaft. 1999.

73. Foissner W, Blatterer H, Berger H. Taxonomische und ökologische revision der ciliaten des saprobiensystems: Band IV: Gymnostomatea, Loxodes, Suctoria. Munich: Informationsberichte des Bayer. . Informationsberichte des Bayerischen Landesamt fur Wasserwirtschaft. 1995.

74. Foissner W, Berger H. A user-friendly guide to the ciliates (Protozoa, Ciliophora) commonly used by hydrobiologists as bioindicators in rivers, lakes, and waste waters, with notes on their ecology. Freshwater Biology. 1996;35(2):375–482.

75. Schwelm A, Berney C, Dixelius C, Bass D, Neuhauser S. The large subunit rDNA sequence of *Plasmodiophora brassicae* does not contain intra-species polymorphism. Protist. 2016;167(6):544–54.

76. Latz MA, Grujcic V, Brugel S, Lycken J, John U, Karlson B, et al. Short-and long-read metabarcoding of the eukaryotic rRNA operon: evaluation of primers and comparison to shotgun metagenomics sequencing. Molecular Ecology Resources. 2022;22(6):2304–18.

77. Callahan BJ, Wong J, Heiner C, Oh S, Theriot CM, Gulati AS, et al. High-throughput amplicon sequencing of the full-length 16S rRNA gene with single-nucleotide resolution. Nucleic Acids Research. 2019;47(18):e103-e.

78. Guillou L, Bachar D, Audic S, Bass D, Berney C, Bittner L, et al. The protist ribosomal reference database (PR2): a catalog of unicellular eukaryote small sub-unit rRNA sequences with curated taxonomy. Nucleic Acids Research. 2012;41(D1):D597–D604.

79. team P. RStudio: Integrated Development Environment for R. Posit Software, PBC, Boston, Massachusetts; 2024.

80. Lu Y, Zhou G, Ewald J, Pang Z, Shiri T, Xia J. MicrobiomeAnalyst 2.0: comprehensive statistical, functional and integrative analysis of microbiome data. Nucleic Acids Research. 2023;51(W1):W310–W8.

81. Oksanen J, Blanchet FG, Friendly M, Kindt R, Legendre P, McGlinn D, et al. vegan: Community Ecology Package. CRAN (Comprehensive R Archive Network); 2007.

82. Ahti L, Sudarshan Shetty, Felix M. Ernst, et al. Orchestrating Microbiome Analysis with Bioconductor [Beta Version] 2021 [Available from: microbiome.github.io/oma/.

